# Identification of smoking-enabled blood miRNA regulatory networks

**DOI:** 10.64898/2026.06.26.733220

**Authors:** Michele Gentili, Brian D. Hobbs, Alla Malinina, Craig P. Hersh, Peter J. Castaldi, Heena Rijhwani, Justin Sui, Corrine Kliment, Michael H. Cho, Kimberly Glass, Enid R. Neptune

## Abstract

Cigarette smoking induces complex signaling disruptions that contribute to diseases such as COPD and lung cancer, yet the molecular mechanisms underlying these effects remain incompletely understood. To address this gap, we analyzed peripheral blood from 3190 COPDGene participants using LIONESS and PUMA and constructed miRNA-mRNA regulatory networks associated with smoking status. Comparing networks for active versus former smokers uncovered a striking shift in regulatory architecture: active smokers exhibited elevated miRNA targeting of the mitochondrial complex I protein *NDUFA12*. This finding was validated in lung tissue expression data from the Lung Genomics Research Consortium (LGRC), where we observed that ever-smokers showed consistent dysregulation of *Ndufa12*-targeting miRNAs compared to never-smokers. This allowed us to identify a set of smoking-defined circulating and tissue-associated miRNAs. To investigate the specific cellular compartment, we analyzed cell-type deconvoluted expression data from COPDGene blood and LTRC (Lung Tissue Research Consortium) lung tissue, as well as lung transcriptomics data from cigarette smoke-exposed mice, and identified the monocyte/macrophage compartment as a principal site of *NDUFA12*/*Ndufa12* expression. Human THP-1 macrophages treated with cigarette smoke extract demonstrated selective inhibition of *NDUFA12* by network-defined miRNAs. These distinct, *NDUFA12*-targeting, smoking-associated miRNA signatures, revealed through network analysis, describe new smoking-mitochondrial interactions that may serve as novel targets for therapeutic intervention.

## Introduction

Smoking represents a systemic insult that functions both as a direct cause of organ-specific diseases (e.g., COPD) and as a powerful risk factor for a broad spectrum of conditions, including solid organ malignancies, stroke, and cardiovascular disease(1, 2). Inhalational exposure to the more than 4,000 constituents of cigarette smoke initiates a complex network of organ- and pathway-specific responses that resist simplification into linear injury cascades(3). Advanced computational platforms offer a powerful way to disentangle this biological complexity, enabling the discovery of mechanistic insights with direct relevance to human health.

MicroRNAs (miRNAs) are ∼22-nucleotide single-stranded RNA molecules that play a pivotal role in the regulation of gene expression by selectively targeting messenger RNAs (mRNAs). Beyond their intracellular functions, miRNAs can be released into the extracellular environment, where they act as intercellular messengers, transmitting signals to both neighboring and distant cells and tissues. The role of miRNAs in mediating smoking-related organ injury is an area of growing scientific interest and investigation. Although several small-scale studies have identified blood-based miRNA panels associated with smoking behaviors, a consistent pattern of target genes or pathways has yet to emerge from large clinical cohorts(4-6). None of these studies generated miRNA and transcriptomic datasets that were later integrated to uncover mechanistically relevant relationships and coordinated biological responses.

Our group has developed two complementary computational tools, PUMA (PANDA Using MicroRNA Associations) and LIONESS (Linear Interpolation to Obtain Network Estimates for Single Samples), that can be combined to infer sample-specific genome wide miRNA-mRNA regulatory networks(7, 8). Regulatory relationships in these networks are not dependent on the fold changes or directionality of responses of individual miRNAs or mRNAs but rather capture the systemic and coordinated responses of sets mRNAs to their upstream miRNA regulators by combining multiple data streams. PUMA (7) is an extension of our message passing framework (PANDA; Passing Attributes between Networks for Data Assimilation) that can be used to refine the characterization of regulatory (e.g. transcription factor or miRNA) engagement with target genes(9). In PUMA, we employ this iterative message-passing approach to integrate miRNA target predictions with gene (mRNA) coexpression data. LIONESS is a mathematical framework that employs a leave-one-out approach to infer sample-specific networks from a population-wide model(8). Given the complexity and clinical relevance of smoking exposure signals that enable injury mechanisms, we used a combination of LIONESS and PUMA to estimate sample-specific miRNA-mRNA regulatory networks in the large well-characterized COPDGene Study and examined the association of these blood miRNA-mRNA relationships with smoking.

We assessed the tissue-context, disease-relevance, and mechanistic impact of the relationships identified using PUMA and LIONESS using (1) lung tissue miRNA-sequencing and gene expression data from the LGRC cohort, (2) chronic CS-exposed murine transcriptomics, and (3) whole cell exposure and genetic targeting studies using a macrophage cell line. Our findings reveal that miRNA modulation of *Ndufa12* expression, a mitochondrial complex I subunit, is linked to smoking status in innate immune cells, suggesting that immune cell energetics reflect the temporal dynamics of adverse exposures.

## Results

### A. Establishment of a miRNA regulatory paradigm by smoking status

We used the LIONESS and PUMA algorithms to estimate subject-specific miRNA-mRNA networks based on blood RNA-sequencing data from 1155 current and 2035 former smokers in the COPDGene Study. Subsequent linear regression analysis identified a miRNA-mRNA regulatory network dysregulated by smoking status (Figure 1A). The identified interactions (edges) between candidate miRNAs and their targeted mRNAs reveal two distinct regulatory programs associated with smoking status; this included 60 significant (FDR<0.05) miRNA-mRNA interactions associated with increased edge weight in current smokers and 20 miRNA-mRNA interactions associated with increased edge weight in former smokers. We calculated the number of significant associated interactions for each gene (mRNA) in these two sets and ranked mRNAs by the difference in the number of associated interactions identified in former versus current smokers. *NDUFA12*, a mitochondrial complex 1 gene, was the top ranked mRNA associated with smoking status and showed regulation by 17 of the 21 miRNAs. We divided these 17 miRNAs into two groups, m1 (n=2) and m2 (n=15), based on whether their regulation was linked to former or current smoking status, respectively. A network visualization of interactions with increased edge weight in current smokers is anchored on *NDUFA12* and its associated m2 miRNAs, but also reveals several genes linked to smoking exposures (PON2, IL1RL1, CLDND1) and mitochondrial functions (SLC35A2, ACOT9, GPAT2)(10-15) (Figure 1B).

**Figure 1.**
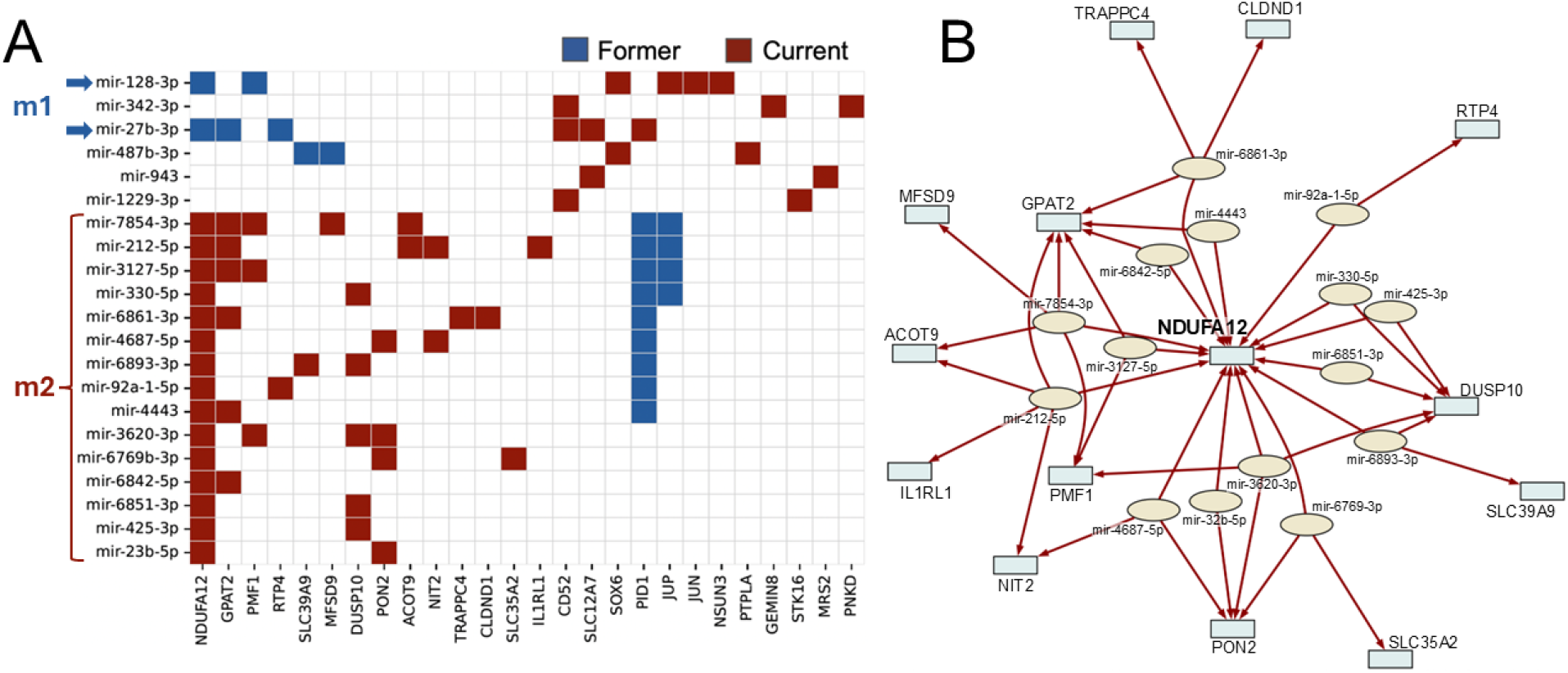
Top ranked miRNA-mRNA interactions by smoking status. (A) A clustered adjacency matrix showing the identified miRNA-mRNA interactions significantly associated with smoking status (FDR<0.05). A filled in square in the matrix represents an identified significant interaction between the miRNA (row) and mRNA (column). The color of the square represents whether the edge has increased weight in former (blue) or current (red) smokers. Within each miRNA and mRNA cluster, rows and columns have been sorted according to the miRNA (row) or mRNA (column) degree (i.e. the total number of significant interactions associated with the miRNA or mRNA). Only miRNAs with a degree of 2 or more are included. (B) A network visualization of the largest connected component of interactions identified as significantly increased in current smokers, showing *NDUFA12* as a central hub in the regulatory network. miRNAs are in pale yellow ovals; target mRNAs are pale blue rectangles.

### B. *NDUFA12* is differentially co-expressed with miRNAs in blood and lung tissue

To assess the context-specific regulation of *NDUFA12* by these 17 miRNAs, we used data from a subset of the COPDGene Study (n=382) for which both mRNA and miRNA expression data were available. We found that both m1 miRNAs had a negative correlation with *NDUFA12* in former smokers, while almost all the m2 miRNAs had a negative correlation with *NDUFA12* in current smokers (Figure 2A). These negative correlations are consistent with a repressive regulatory role of these miRNAs on *NDUFA12* expression, as inferred by the LIONESS-PUMA network analysis, for which we did not use the miRNA expression.

**Figure 2.**
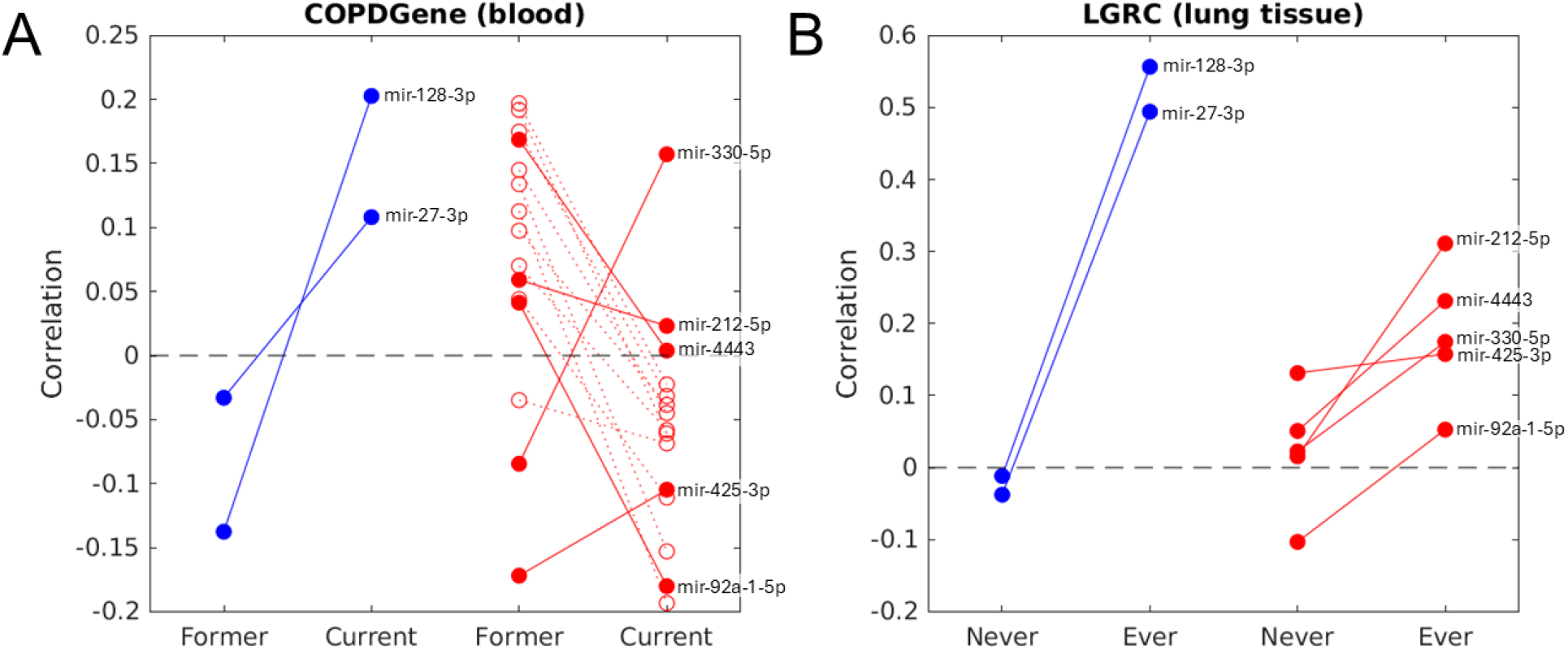
Characterization of smoking status subgroups of NDUFA12-regulating miRNAs by correlation with *NDUFA12* mRNA in COPDGene and LGRC. Correlation values of network-identified miRNAs with *NDUFA12* in (A) COPDGene (blood) and (B) LGRC (lung tissue) when samples in the datasets are stratified based on smoking status. miRNAs were assessed in two groups, m1 (blue) and m2 (red), based on whether their regulation of *NDUFA12* was linked to former or current smoking status, respectively (see Figure 1). miRNAs with expression data in both datasets are shown with solid circles and lines, the subset of m2 miRNAs that did not have measurable expression in LGRC but were detected in COPDGene are shown with hollow circles and dashed lines. miRNAs expressed in both COPDGene and LGRC are labeled.

We further evaluated the tissue-context of these interactions using data from lung tissue samples from the Lung Genome Research Consortium (LGRC). We leveraged 351 samples for which both mRNA expression and miRNA sequencing data was available. This dataset contained expression values for a subset (7 out of 17) of the miRNAs targeting *NDUFA1*, including both m1 miRNAs and 5 of the m2 miRNAs. In addition, rather than former vs current smoking status, as in the COPDGene Study, the smoking status of LGRC subjects were categorized as never vs ever smokers. Despite these differences, we observed the same trend for both m1 miRNAs in LGRC as we observed in COPDGene (Figure 2B). Looking at the individual miRNA changes in the co-expression with *NDUFA12* (Table 1) we found mir-128 (m1), mir-27b-3p (m1), mir-330-5p (m2), and mir-425-3p (m2) have similar directional behaviors in both datasets, while miR-212-5p (m2), mir-92a-1-5p (m2), and mir-4443 (m2) have opposing patterns. We termed these miRNAs concordant versus discordant, respectively.

**Table 1.** Correlation of miRNAs in m1 and m2 with *NDUFA12* by smoking status in both COPDGene and LGRC. Concordant: Change in miRNA correlation with NDUFA12 (ΔCorr) in former vs current (blood) and never vs ever (lung tissue) has the same sign. Discordant: Change in miRNA correlation with NDUFA12 (ΔCorr) in former vs current (blood) and never vs ever (lung tissue) has the opposite sign.

|  |  | COPDGene (whole blood) |  |  | LGRC (lung tissue) |  |  |  |
| --- | --- | --- | --- | --- | --- | --- | --- | --- |
| Group | miRNA | Smoking Status | Corr | $\Delta$ Corr<br>(former-current) | Smoking Status | Corr | $\Delta$ Corr<br>(never-ever) | Concordant? |
| m1 | mir-128-3p | former | -0.137587 | -0.340363 | never | -0.011692 | -0.568252 | concordant |
|  |  | current | 0.202776 |  | ever | 0.55656 |  |  |
|  | mir-27b-3p | former | -0.032945 | -0.141079 | never | -0.037623 | -0.531794 | concordant |
|  |  | current | 0.108134 |  | ever | 0.494171 |  |  |
| m2 | mir-212-5p | former | 0.059195 | 0.036054 | never | 0.015527 | -0.295662 | discordant |
|  |  | current | 0.023141 |  | ever | 0.311189 |  |  |
|  | mir-330-5p | former | -0.084492 | -0.241749 | never | 0.022471 | -0.152054 | concordant |
|  |  | current | 0.157257 |  | ever | 0.174525 |  |  |
|  | mir-92a-1-5p | former | 0.041309 | 0.221454 | never | -0.10321 | -0.155986 | discordant |
|  |  | current | -0.180145 |  | ever | 0.052776 |  |  |
|  | mir-4443 | former | 0.168592 | 0.16472 | never | 0.05094 | -0.180279 | discordant |
|  |  | current | 0.003872 |  | ever | 0.231219 |  |  |
|  | mir-425-3p | former | -0.17192 | -0.067117 | never | 0.131278 | -0.026389 | concordant |
|  |  | current | -0.104803 |  | ever | 0.157667 |  |  |

We used miRTarBase, a web tool incorporating levels of experimental validation, to identify other complex I proteins that show evidence of regulation by these seven miRNAs. Although *NDUFA12* was not identified, mitochondrial complex I genes are plausible targets for five of these miRNAs (Supplemental Table 1). Mitochondrial disturbances have been reported for all four of the concordant miRNAs but only one of the discordant miRNAs. Association with smoking exposure has been shown for two of the concordant and one of the discordant miRNAs.

### C. *NDUFA12* expression in alveolar macrophages associates with smoking status

We next assessed *NDUFA12* expression in specific blood and lung cell types using deconvoluted expression data from COPDGene(16) and LTRC(17), respectively. In whole blood, *NDUFA12* was expressed in many immune cell types, including mast cells, CD4 and CD8 T cells, monocytes, and B cells (Supplemental Table 2). In lung tissue *NDUFA12* was expressed in mast cells, vascular endothelial cells, goblet cells, alveolar macrophages, and other cell types (Supplemental Table 3). We determined whether smoking status was associated with *NDUFA12* expression levels in specific lung cell types and found a suggestive association (pvalue=0.07) of *NDUFA12* expression in alveolar macrophages by ever versus never smoking status after adjusting for gender, age, BMI, and FEV_1_/FVC (Table 2). Of note, removing the FEV_1_/FVC covariate decreased the P-value to 0.03.

**Table 2.** Coefficients from the linear model evaluating the differential expression of *NDUFA12*. Based on deconvoluted alveolar macrophage expression data from TOPMed LTRC.

| Model | ~ smoking + sex + age + bmi + fev1fvc |  |  |  | ~ smoking + sex + age + bmi (no fev1fvc) |  |  |  |
| --- | --- | --- | --- | --- | --- | --- | --- | --- |
| Variable | coef<br>[0.025, 0.975] | std err | t | pvalue | coef<br>[0.025, 0.975] | std err | t | pvalue |
| intercept | 0.2263<br>[0.012, 0.44] | 0.109 | 2.075 | 0.0384 | 0.259<br>[0.052, 0.466] | 0.105 | 2.463 | 0.0141 |
| smoking-status | -0.2071<br>[-0.433, 0.018] | 0.115 | -1.804 | 0.0717 | -0.2411<br>[-0.459, -0.023] | 0.111 | -2.176 | 0.0300 |
| sex | -0.0899<br>[-0.254, 0.074] | 0.083 | -1.078 | 0.2816 | -0.1006<br>[-0.263, 0.062] | 0.083 | -1.212 | 0.2258 |
| age | -0.016<br>[-0.095, 0.063] | 0.04 | -0.397 | 0.6916 | -0.0151<br>[-0.094, 0.064] | 0.04 | -0.375 | 0.7080 |
| BMI | -0.0256<br>[-0.11, 0.058] | 0.043 | -0.6 | 0.5489 | -0.0116<br>[-0.092, 0.069] | 0.041 | -0.283 | 0.7770 |
| FEV1/FVC<br>(post) | 0.0492<br>[-0.036, 0.135] | 0.043 | 1.131 | 0.2583 | NA | NA | NA | NA |

### D. Expression of *NDUFA12* and other complex I components in murine chronic smoke exposure model

Current versus former smoking status is challenging to dissect in a murine model in which the time frames of relevant exposure are significantly shortened and the definition of former status is unresolved. However, the ever versus never smoking status qualified in LGRC (Figure 2B) could be reasonably simulated with the chronic CS-exposed mouse model. Consequently, we examined expression of *Ndufa12* in a single cell RNA sequencing data set from the lungs of mice with chronic CS exposure for 6 months (GSE277533)(18, 19). *Ndufa12* expression appeared higher in alveolar macrophages in CS-exposed compared to air-exposed control mice (Supplemental Figure 1). Since *Ndufa12* resides in the N-module of the mitochondrial Complex I, we examined the expression and CS-mediated dysregulation of other proteins within this module in alveolar macrophage subtypes. Across these cell types, selective module components were differentially expressed in alveolar macrophage compartments with nominally significant changes in *Ndufa12, Ndufa7, Ndufs7*, and *Ndufv1* expression (Supplemental Table 4).

### E. *Ndufa12*-targeting miRNAs in THP-1 cells exposed to CS

We exposed the human macrophage cell line THP-1 to cigarette smoke extract and measured the effect of five miRNAs (two in m1 and three in m2) on *NDUFA12* expression (Figure 3A). The selected miRNAs were those identified as having evidence for targeting mitochondrial complex I genes (see Supplemental Table 1). While mimics of two of the three m2 miRNAs, miR-92a and miR-425, showed CS specific downregulation of *NDUFA12*, mimics of both m1 miRNAs, miR-27b and miR-128, did not (Figure 3B). Mimics of miR-330-5p (m2) downregulated *NDUFA12* with and without CS exposure but showed more robust effects with concurrent CS treatment. Overall, the correlation of m2 miRNAs, but not m1 miRNAs, with smoking behaviors suggest CS-enhanced regulation of *NDUFA12* expression. Additionally, discordant vs concordant miRNA assignments were not strictly associated with miRNA mimic responses.

**Figure 3.**
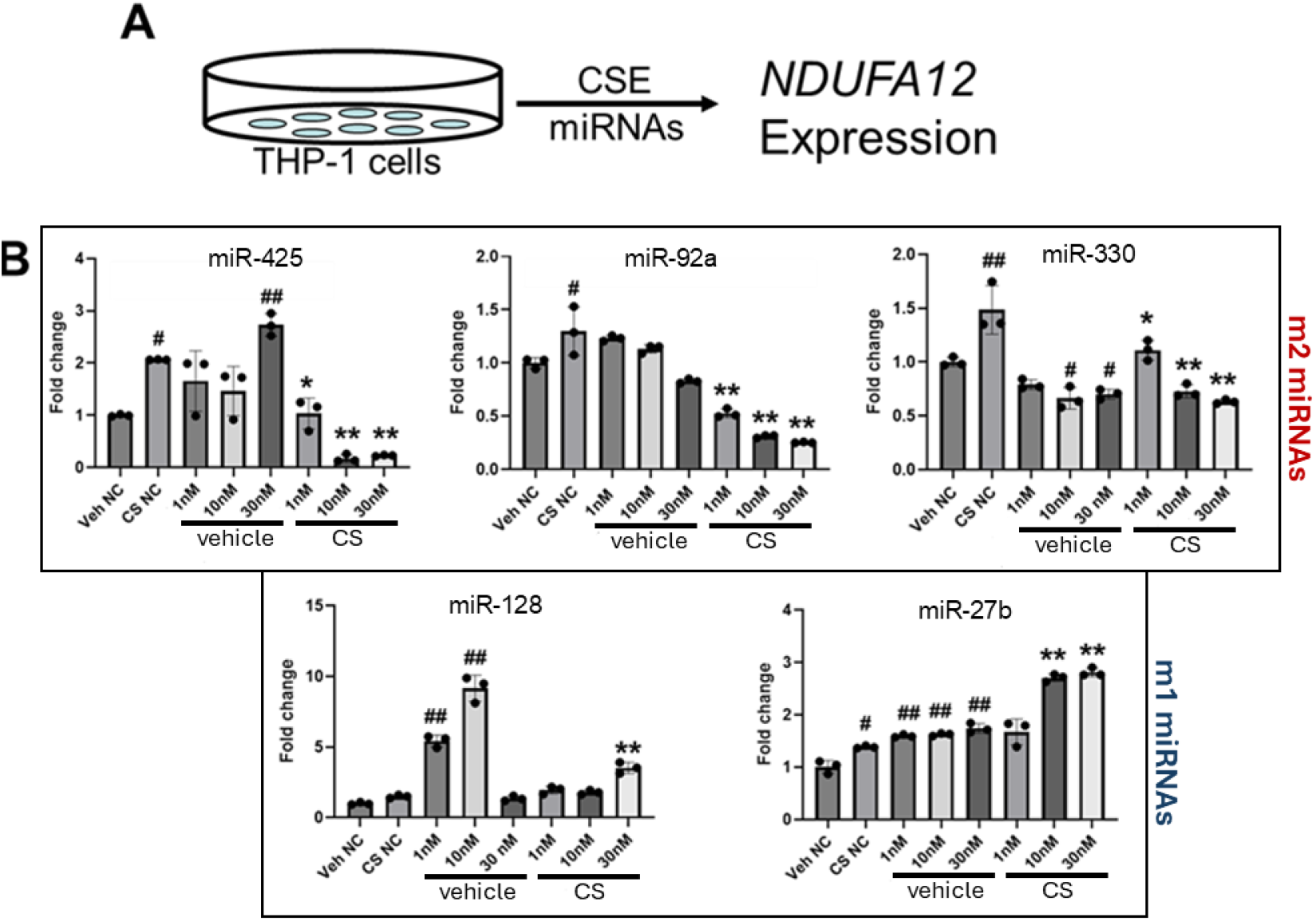
Experimental assessment of m1 and m2 miRNA modulation of *NDUFA12* expression. (A) Schematic of miRNA administration to CSE-exposed THP-1 cells. (B) Effect of miRNA mimic dosage on NDUFA12 expression. Expression was measured by qPCR after 8h treatment with mimics followed by 24h exposure to CS/CSE (cigarette smoke extract) or vehicle (Veh). Fold change is NDUFA12 expression normalized to GAPDH. miRNAs were assessed in two groups, m1 (bottom) and m2 (top), based on whether their regulation of NDUFA12 was linked to former or current smoking status, respectively (see Figure 1). #p<0.05, ##p<.001 compared with Veh NC. *p<0.05, **p<.001 compared with CS NC. NC-Scrambled control. All studies performed in triplicate.

## Discussion

We used an advanced bioinformatics and network modeling approach to identify candidate biomarkers of current smoking exposure. Networks modeled using blood expression data from COPDGene Study showed a relationship between active smoking with miRNA regulation of the complex I mitochondrial gene *NDUFA12*. Analysis of expression data collected in lung tissue samples from never- and ever-smokers confirmed a smoking associated relationship between selected miRNAs and *NDUFA12* expression. Expression of *NdufA12* in subtypes of alveolar macrophages with chronic CS exposure in a murine model clarified compartmental effects of smoking behaviors which was further supported by CS treatment of a macrophage cell line. As such, we demonstrate an active smoking-enabled network of miRNA governed mitochondrial dysregulation that converges onto the innate immune compartment.

Circulating cell-free microRNAs (miRNAs) have emerged as a novel tool for biomarker discovery with high translational relevance, especially for neoplastic disorders(20, 21). Several recent studies report miRNA panels in lung, lung cells, or blood that associate with smoking behaviors(4-6, 22). Limitations of these studies, even when conflated with target RNAs by sequence or coexpression, include small cohort size, use of predefined exploratory panels, use of array platforms and a lack of methodologic validation across multiple target prediction resources and tissue types. Our application of the PUMA and LIONESS algorithms and our use of two distinct sample sources alleviates many of these concerns. Despite these differences, three of the miRNAs identified in our COPDGene and LGRC analysis, miR-92a, mir-27b and miR-128, were also detected in other profiles of smoking behaviors (4, 5, 22-24). Our validation using the THP-1 cell line showed that of these three molecules only miR-92a displayed *NdufA12* modulation by cigarette smoke extract exposure.

Willinger et al. conducted a whole blood miRNA and mRNA profiling study of ∼5000 Framingham Heart Study participants defined by smoking status(6). They employed a high throughput qPCR approach (detectable miRNAs) and a human microarray gene chip (mRNAs). Upon applying an FDR threshold, only five miRNAs were differentially expressed in current versus former smokers with only one showing an upregulation. The assignment of these 5 miRNAs with coexpressed predicted target genes did show GO enrichment for T-cell homeostasis. None of these miRNAs or highly ranked target mRNAs were identified in our study as associated with current smoking status. This may reflect differences in whole blood vs cell free sampling or the overrepresentation of persons with lung disease in the COPDGene and LGRC populations.

Systemically, active smoking reflects a chronic, cyclic, and sustained injury process whereas former smoking represents a low grade inflammatory injury that degrades over time but never fully resolves(25). Further, acute effects of nicotine exposure, especially in brain function, are only evident with active smoking(26). The systemic component of these states is of interest given the multiorgan consequences of these exposures and the only partial mitigation of disease risk with smoking cessation. The set of miRNAs we identified that had increased regulatory targeting in the context of active smoking in COPDGene includes regulators of malignancy progression(27-32), inflammation(33-36), vascular development/remodeling(37-40), and COPD(41-43), all processes associated with smoking enabled disease states. As such, this set of miRNA regulators reflects the breadth of smoking related pathologies evident in end organ disease.

Most clinical and preclinical assessments of mitochondrial injury with smoking focus on specific disease outcomes, COPD or malignancy, rather than smoking behaviors as the informative phenotypes. Our study provides a possible explanation for chronic cigarette smoke-induced alterations in macrophage energetics— manifested as compromised host responses, lipid dysregulation, reduced pathogen clearance, and impaired oxidative stress management—by invoking upstream miRNA regulatory drivers(44-48). One compelling finding of a small microarray profiling study of alveolar macrophages in smokers is a broad downregulation of miRNA expression across several pathway domains, a readout not explored in our analysis(49). Using blood data from COPDGene, we show that active smoking compared to former smoking dramatically increases multiple regulatory miRNA-mRNA interactions that converge on *NDUFA12*. Our subsequent analyses in bulk lung tissue data and cell-type specific data suggest that the balance of circulating versus lung-expressed miRNAs targeting *NDUFA12* show both concordant and discordant alignments that may reflect nuances of smoking behaviors. Additionally, lung function may be a mediator of the effect of smoking status with miRNA-governed *NDUFA12* expression.

NDUFA12 is a protein component of the N subdomain of mitochondrial complex I and operates within a cooperative multiprotein cluster to form a proton gradient for ATP generation(50). Alterations in the abundance of any component of Complex I causes instability of the structure compromising the efficient generation of ATP. Loss-of function variants in *NDUFA12* cause Leigh Syndrome, a rare progressive neurodegenerative disorder of mitochondrial origin which develops during early postnatal life(51, 52). Defects in complex I reductase activity associate with smoking in platelets but are unexamined in other immune cell types(53). Wain also identified NDUFA12 as a possible druggable target for lung function maintenance using a large COPD GWAS study(54); notably, genetic evidence, including from GWAS, increases the probably of success in a clinical trial by over 2-fold(55). Our survey of *NdufA12* and other nuclear mitochondrial complex I genes in a murine chronic smoke exposure model established a strong dysregulation in the alveolar macrophage compartment consistent with broad complex l impairment.

Our study has notable strengths. The human cohorts we leverage are large and well characterized. Both COPDGene and LGRC use a sequencing approach for miRNA profiling which recovers a broader range of molecules compared to chip-array strategies. We also employ complementary discovery and validation datasets which extend our systemic findings to organ specific signaling disturbances conferred by smoking behaviors. Finally, we use a preclinical murine CS exposure platform and whole cell studies to refine seminal aspects of the smoking-miRNA-mRNA paradigm that we advance here. Limitations are also evident. In the human cohorts we use both persons with and without chronic lung disease such as COPD and interstitial lung disease. Smoking status was also defined differently in the two studies, dichotomized as current versus former in COPDGene and ever versus never in LGRC. The definitive source of the miRNAs in the lung and blood samples also cannot be fully specified despite our analysis of deconvoluted gene expression data.

The role of extracellular miRNAs in tissue maintenance and systemic homeostasis is widely debated. Whether these miRNAs reflect cell destruction or a means of directive cell-cell communication, e.g. exosomes, is unresolved. A mouse survey of miRNAs in different organs and body fluids showed a sizable contribution of lung-sourced miRNAs in blood profiles(56). Our approach, with a hierarchical integration of advanced bioinformatics methods and data from multiple human and murine sources, coupled with a systemic toxic exposure, provides a powerful way to reconcile features of exposure (such as current versus former smoking) with organ-specific injury responses. Moreover, our identification of a smoking-dependent miRNA network that targets a component of the mitochondrial machinery invites further exploration of computational approaches to parsing exposome-epigenetic engagements.

## Methods

### Bulk mRNA and miRNA expression data for blood (COPDGene) and lung tissue (LTRC/LGRC)

*<u>COPDGene</u>* (clinicaltrial.gov identifier: NCT00608764) is a multicenter longitudinal observational study designed to identify genetic factors associated with COPD. Further details regarding recruitment have been previously published(57). All participants provided written, informed consent and institutional review boards approved the study at all participating centers.

In our analysis, we leveraged previously generated blood RNA-sequencing and blood miRNA-sequencing data(58) from COPDGene, processed as described in Gentili et al.(59). For the RNA-sequencing data, we analyzed phase 2 data, removing batches with less than 10 subjects, PRISm subjects, and never smokers, resulting in data from 3190 subjects. For miRNA-sequencing data, we analyzed phase 2 data from 439 subjects to identify expressed miRNAs, yielding 679 potential regulatory species.

<u>*The Lung Tissue Research Consortium (LTRC)*</u>, funded by the National Heart, Lung, and Blood Institute (NHLBI), collected lung tissue from donors undergoing clinically indicated thoracic surgery. In this work, we analyzed data from two subsets of LTRC.

The first, the *LGRC* (Lung Genomics Research Consortium) was a multi-institutional program that generated genomic data for a subset of LTRC samples, to better understand the genetic and molecular mechanisms underlying chronic lung diseases, including COPD and ILD. In our analysis, we leveraged previously generated microarray gene (mRNA) expression data (GSE47460)(60, 61) and miRNA-sequencing expression data (GSE201121)(62) from LGRC. The microarray gene expression data was processed as previously described(63, 64). Briefly, because these data were collected using two array platforms, each array was first separately background corrected and rma-normalized(65). Then, to merge the platforms, duplicate probes were removed and any remaining non-identical probes were matched between the platforms based on gene name. The merged data was subsequently corrected for array-specific batch effects using ComBat(66). For the miRNA-sequencing data, normalized miRNA expression data was directly downloaded from the gene expression omnibus GSE201121. In total, we identified 351 subjects in LGRC that had both gene (mRNA) expression and miRNA-sequencing expression data.

The second LTRC dataset (referred to as simply LTRC, phs001662) is part of the Trans-Omics in Precision Medicine (TOPMed) study, a multi-center, cohort, and omics study. Approximately 1500 TOPMed samples from individuals with COPD, ILD, and without chronic lung disease (controls), selected independently from LGRC, underwent multiomic profiling, including RNA-sequencing. The LTRC RNA-sequencing data has been deconvoluted to identify lung cell-type specific gene expression (see below).

### COPDGene and LTRC cell-type specific expression based on deconvolution of RNA-sequencing data

To interrogate cell-type specific expression information, we leveraged cell-type specific deconvoluted gene expression from Ryu et al.(16, 17). Data were generated by applying CIBERSORTx to (1) LTRC RNA-sequencing data using the Adams et al. (lung) reference panel (67) and (2) COPDGene RNA-sequencing samples using the LM22 (blood0) reference panel(68).

### Clinical Variables

In COPDGene, current smokers were those who reported smoking in the past month at the time of the Phase 2 blood draw; the remainder of subjects were former smokers. For the LGRC data, data reported ever or never smoker status; an ever smoker was defined as > 100 lifetime cigarettes.

### LIONESS-PUMA

We created sample-specific miRNA-mRNA regulatory networks for 3190 subjects in COPDGene as described in Gentili et al.(59). Briefly, we downloaded miRNA targeting information from TargetScan. We then filtered this targeting information to only retain miRNAs and mRNAs expressed in the COPDGene miRNA-sequencing and RNA-sequencing data, respectively. This resulted in a miRNA-mRNA map between 570 miRNAs and 11859 mRNAs with measurable expression in the COPDGene RNA-sequencing data, and a final gene expression dataset containing 3190 subjects and 11,859 mRNAs (genes).

Using our dataset of 3190 subjects and 11,859 mRNAs, we created sample-specific mRNA correlation networks using LIONESS. We then applied PUMA to each of these sample-specific correlation networks to integrate them with miRNA-mRNA targeting information, creating sample-specific miRNA-mRNA regulatory networks. Note that miRNA expression information was only used to identify which miRNAs to model using PUMA, but not for the miRNA-mRNA network reconstruction. For more information on the construction of these networks, see Gentili et al.(59).

### Covariate Adjusted Network Analysis

We used the *statsmodels* python package (v 0.14.2) to perform linear regression on the LIONESS-PUMA-estimated sample-specific miRNA-mRNA interaction scores with the following model:

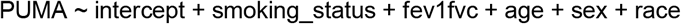

We identified miRNA-mRNA interactions whose edge weight changed significantly (FDR <0.05, Benjamini/Hochberg) as a function of smoking status, adjusting for FEV_1_/FVC ratio, age, sex, and race. Similar to the process described in Gentili et al.(59), we identified a ‘core’ network among the identified edges that included miRNAs with at least two significant interactions and their target mRNAs.

### Mouse chronic cigarette smoke (CS) exposure transcriptomics

Previously published lung transcriptomes generated from C57BL/6 mice exposed to 6 months of mainstream CS compared to room air controls were downloaded from the Gene Expression Omnibus (GSE277533) and examined for differential expression of mitochondrial complex 1 N module genes (19). Cell type assignments employed published annotations(69). For details, see supplemental methods.

### Whole Cell Validation

THP-1 cells, a human leukemia cell line, were purchased from the ATCC, and cultured in DMEM with 10% FBS containing 100 U/ml penicillin and 100 μ g/ml streptomycin. The cells were plated into 6-well dishes and incubated in a humidified incubator with an atmosphere of 5% CO2 at 37° C. The interval of cell passage was nearly 48h, and the 5th to 15th passages of cells were used in this experiment. CS extract was added in different concentrations and harvested at 24 and 48h after exposure. Cells were harvested and RNA recovered with Trizol.

### Whole Cell qPCR Analyses

Total RNA isolated from cells with Trizol-based protocol was treated with DNase and reverse-transcribed as described. The expression levels of target genes were determined in triplicate or quadruplicate from the standard curve and normalized to the Gapdh mRNA level. Probes were obtained from Applied Biosystems (ThermoFisher, Waltham, MA).

### MiRNA transfection

MiRNA mimics for *mir-128, -27b, -425, -92a*, and *-330-5p* were obtained from ThermoScientific (Waltham, MA) and delivered to cells. THP-1 cells were harvested and RNA recovered with Trizol as described.

### Statistics

For miRNA mimic studies, comparisons between more than two groups were performed by one way analysis of variance (ANOVA), whereas comparisons with two or more independent variable factors were performed by two-way ANOVA using GraphPad Prism 9.0 software. Statistical tests used biological replicates with p<0.05 were considered statistically significant.

## Author Contributions

MG, BH, MC, KG, and ERN conceived, coordinated and designed the study. BH, HR, JS, CK, MC, and KG retrieved the datasets and did the analysis. AM and JS performed the experiments. MC, KG, and ERN wrote the manuscript. MG, BH, AM, CH, PC, and CK critically reviewed and edited the manuscript. All authors have reviewed and approved the final manuscript before submission.

## Acknowledgements

Molecular data for the Trans-Omics in Precision Medicine (TOPMed) program was supported by the National Heart, Lung and Blood Institute (NHLBI).

