## Supplemental Materials and Methods for "Identification of smoking-enabled blood miRNA regulatory networks"

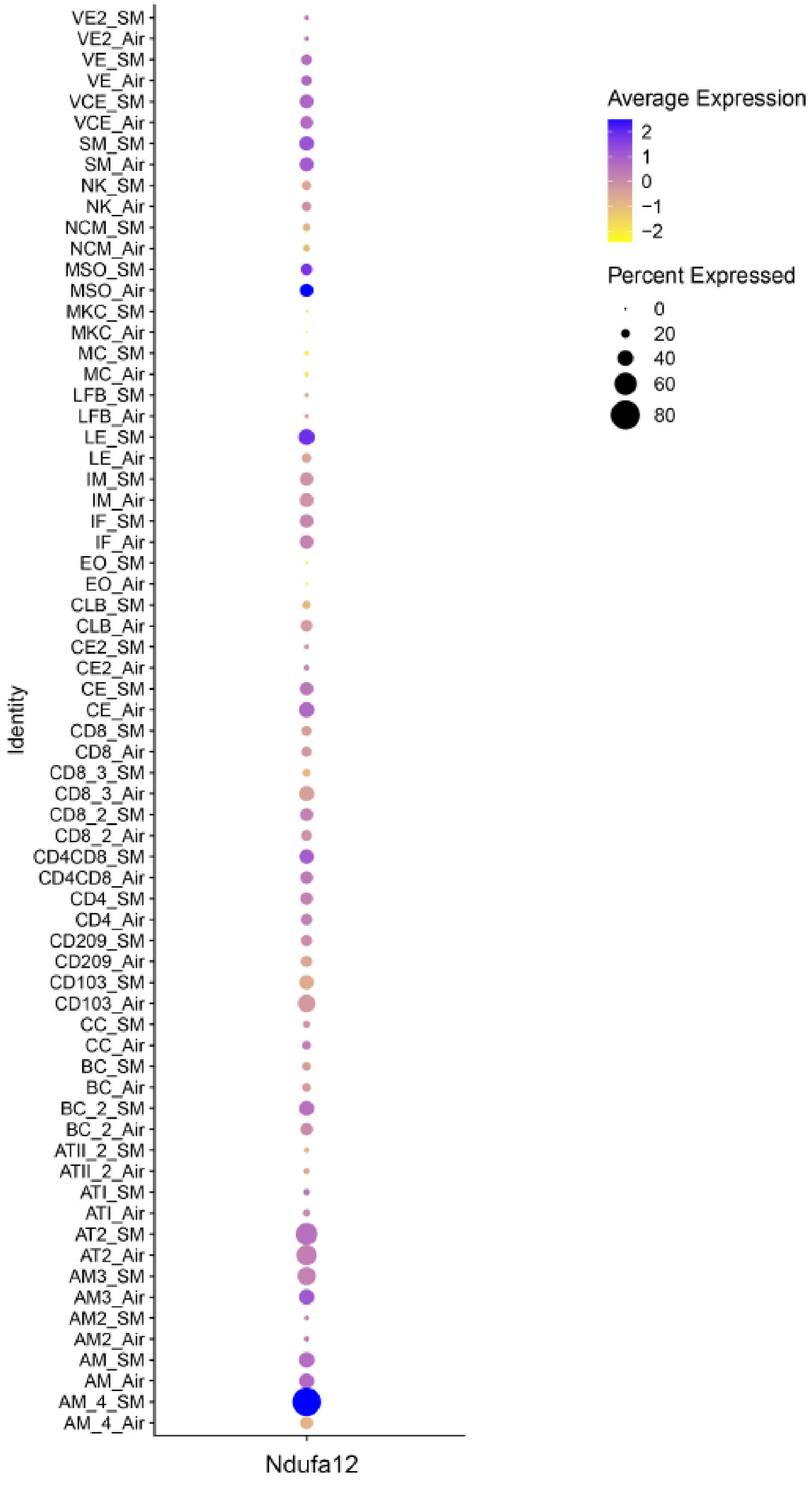

**Supplemental Figure 1: Single-cell RNA-sequencing of cigarette smoke exposed mouse lung for Ndufa12 expression.** *Ndufa12* expression in single-cell RNA-sequencing data (GSE277533) from C57BL/6 mice exposed to air or cigarette smoke for 6 months (n=4 mice per treatment group, individually sequenced). Circle size represents the percentage of cells expressing the gene and color gradient represents the average expression of cells. Cell identities include VE – vascular endothelial, VCE – VCAM+ endothelial, SM – smooth muscle, NK – natural killer, NCM – non-classical monocytes, MSO – mesothelial, MKC – megakaryocyte, MC – mast cells, LFB – lipofibroblasts, LE – lymphatic endothelial, IM – interstitial macrophage, IF – interstitial fibroblast, EO – eosinophil, CLB – club cell, CE – capillary endothelial, CC – ciliated cell, BC – basal cell, ATII – alveolar type 2, ATI – Alveolar type 1, AM – alveolar macrophage.

| COPDGene:LGRC miRNA-NDUFA12 concordance | MicroRNA (m1/m2) | Mitochondria | Smoking | Complex I Target Genes (miRTARBase) | References |
|---|---|---|---|---|---|
| Concordant | miR-128 (m1) | Yes | Yes | NDUFS5<br>NDUFC2<br>Ndufs4 | 26010876,<br>34384932,<br>38422767,<br>22512273 |
| | miR-27b (m1) | Yes | Yes | NDUFA10<br>NDUFA5<br>NDUFC2<br>NDUFS6<br>Ndufa2<br>Ndufs4<br>Ndufs6 | 28698281,<br>26452502,<br>27456084,<br>33671744,<br>39616277 |
| | miR-330-5p (m2) | Yes | NR | NDUFA2 | 32360974,<br>32392088 |
| | miR-425-3p (m2) | NR | NR | NDUFA7 | |
| Discordant | mir-212-5p (m2) | NR | NR | NR | |
| | miR-4443 (m2) | NR | NR | NR | |
| | miR-92a (m2) | Yes | Yes | NDUFA5<br>NDUFA7<br>NDUFB10<br>NDUFC2<br>NDUFV3<br>Ndufa11<br>Ndufa5 | 31319246,<br>38173217,<br>31692088,<br>34097306 |

**Supplemental Table 1: miRTarBase information for miRNAs targeting NDUFA12, separated based on whether they have concordant or discordant differential correlation patterns with smoking in COPDGene versus LTRC.** Target genes show experimental validation by platform criteria. Mitochondria and Smoking responses indicate published reports associating the miRNA with the compartment or the exposure, respectively. Red denotes N-module genes. NR-Not reported. Information from https://mirtarbase.cuhk.edu.cn

| **Cell Type** | **Average NDUFA12 Expression** |
|---|---|
| Mast cells | 106.52645 |
| T cells CD8 | 38.056032 |
| T cells CD4 | 31.054449 |
| Monocytes | 21.506796 |
| B cells | 18.326294 |
| Neutrophils | 1.000000 |
| Eosinophils | 1.000000 |
| Dendritic cells | 1.000000 |
| NK cells | 1.000000 |
| Plasma cells | 1.000000 |

**Supplemental Table 2: Cell-type specific expression of *NDUFA12* in blood.** Average expression of *NDUFA12* in various cell types based on cell-type specific deconvoluted gene expression data from blood samples in COPDGene.

| **Cell Type** | **Average *NDUFA12* Expression** |
|---|---|
| Mast | 81.306376 |
| ILC A | 75.91499 |
| VE Capillary A | 41.537778 |
| VE Capillary B | 34.459278 |
| Goblet | 34.369757 |
| Macrophage Alveolar | 33.775378 |
| T Cytotoxic | 31.886571 |
| SMC | 30.81322 |
| Fibroblast | 25.755755 |
| Club | 22.987959 |
| Aberrant Basaloid | 21.396707 |
| ATI | 18.961717 |
| Macrophage | 17.528997 |
| ATII | 15.060108 |
| Myofibroblast | 10.348839 |
| B Plasma | 9.159881 |

**Supplemental Table 3: Cell-type specific expression of *NDUFA12* in lung tissue.** Average expression of *NDUFA12* in various cell types based on cell-type specific deconvoluted gene expression data from lung tissue samples in LTRC. VE = vascular endothelial.

| | Unadj p-value | FC |
|---|---|---|
| **AM1 (1076)** | | |
| ***Ndufs7**** | 0.048 | -0.127 |
| **AM2 (1063)** | | |
| ***NdufA7**** | 0.022 | 0.237 |
| **AM3 (381)** | | |
| ***NdufA7**** | 0.005 | 0.746 |
| ***NdufV1**** | 0.027 | 0.779 |
| ***NdufS2*** | 0.066 | 0.523 |
| **AM4 (166)** | | |
| ***NdufA12**** | 0.026 | 1.03 |
| ***NdufA7**** | 0.001 | 1.104 |

**Supplemental Table 4: Differential Expression of Mitochondrial Complex 1 N-Module Genes in Murine Alveolar Macrophage Sub-populations by CS exposure.** AM1-4 – Alveolar macrophage subclusters 1-4 (see Supplemental Methods). Number of cells annotated to each cell type is noted in parentheses. Unadj p-value – Unadjusted p-value. FC – fold change (smoke vs room air). * and red font – nominally significant differential expression.

### Supplemental Methods

#### Single cell RNA sequencing (scRNAseq) data analysis and clustering

scRNA seq data from C57BL/6 mice exposed to 6 months of mainstream CS compared to room air controls (GSE277533)[1, 2] were analyzed for differential expression of mitochondrial complex 1 N module genes. Animals in this data set were exposed to 4 unfiltered cigarettes/day, 5 days/week for 6 months (University of Kentucky) according to previously published protocols [3, 4]. The analysis of the scRNA sequencing data was conducted with Seurat V4.1.0 R package and R V4.0.2 [5, 6]. Cells were screened according to the following criteria: greater than 200 genes, fewer than 3000 genes, and less than 25% mitochondrial genes. Following the standard Seurat workflow, the data was normalized, and clustering and visualization were performed using the uniform manifold approximation and projection (UMAP) method. To assign cell identities, canonical cell markers and gene marker lists established previously were used[7]. Mitochondrial complex I genes were first analyzed across all cell types (Supplemental Figure 1); this included four alveolar macrophage (AM) clusters with sufficient cells for differential expression analysis (number of cells ≥150). To achieve greater precision for macrophages, the alveolar macrophage (AM) clusters were combined and re-clustered. Within each of the subclustered AMs with sufficient cells (≥150), differential expression of mitochondrial complex I genes was analyzed using the *FindMarkers* Seurat function (Supplemental Table 4).
